## Supplementary material for "Towards a comprehensive view of the pocketome universe – biological implications and algorithmic challenges": S1Table.docx

**S1 Table.** **Additional information for all predicted pockets including the number of FoldSeek (FS) cluster per species, the ratio of pockets per protein, the number of pockets before filtering, and the number of pockets that did not pass the respective quality check.**

| **Org.** | **Gene dupl. [%]** | **# pockets before filtering** | **# pLDDT < 70** | **# PAE > 10** | **# pockets / # proteins** | **# FS cluster** |
| --- | --- | --- | --- | --- | --- | --- |
| ECOLI | 0.9 % | 2,764 | 82 | 102 | 0.694 | 3,010 |
| YEAST | 2.2 % | 3,596 | 497 | 535 | 0.534 | 4,730 |
| CANAL | 12.6 % | 3,573 | 334 | 400 | 0.556 | 4,930 |
| ARATH | 34.7 % | 12,460 | 971 | 1,209 | 0.447 | 14,325 |
| ORYSJ | 6.3 % | 14,571 | 3,493 | 3,637 | 0.346 | 21,379 |
| MAIZE | 53.3 % | 16,941 | 3,020 | 3,301 | 0.375 | 20,896 |
| SOYBN | 73.6 % | 24,372 | 2,599 | 2,943 | 0.44 | 20,476 |
| DROME | 58.2 % | 6,425 | 345 | 478 | 0.474 | 9,652 |
| CAEEL | 25.7 % | 11,439 | 2,224 | 2,243 | 0.497 | 11,794 |
| MOUSE | 48.0 % | 11,023 | 438 | 785 | 0.498 | 13,020 |
| HUMAN | 62.4 % | 9,016 | 569 | 816 | 0.421 | 13,424 |
