## Supplementary material for "Towards a comprehensive view of the pocketome universe – biological implications and algorithmic challenges": S2Table.docx

**S2 Table.** **Number of unique pockets, singletons, and real clusters with threshold 5 applied to the ProBiS alignment score.**

| **Species** | **# Unique BS** | **# Singletons** | **# Real clusters** |
| --- | --- | --- | --- |
| ECOLI | 1,488 | 1,208 | 280 |
| YEAST | 1,639 | 1,291 | 348 |
| CANAL | 1,713 | 1,397 | 316 |
| ARATH | 2,529 | 1,692 | 837 |
| ORYSJ | 2,703 | 1,883 | 820 |
| MAIZE | 3,019 | 2,054 | 965 |
| SOYBN | 3,265 | 1,889 | 1,376 |
| DROME | 2,453 | 1,939 | 514 |
| CAEEL | 3,178 | 2,544 | 634 |
| MOUSE | 2,690 | 1,982 | 708 |
| HUMAN | 2,595 | 1,877 | 718 |

The threshold was applied to the similarity matrix that was obtained using the alignment scores of ProBiS.
