## Supplementary figures and images for "Towards a comprehensive view of the pocketome universe – biological implications and algorithmic challenges"

### S1Fig.tiff

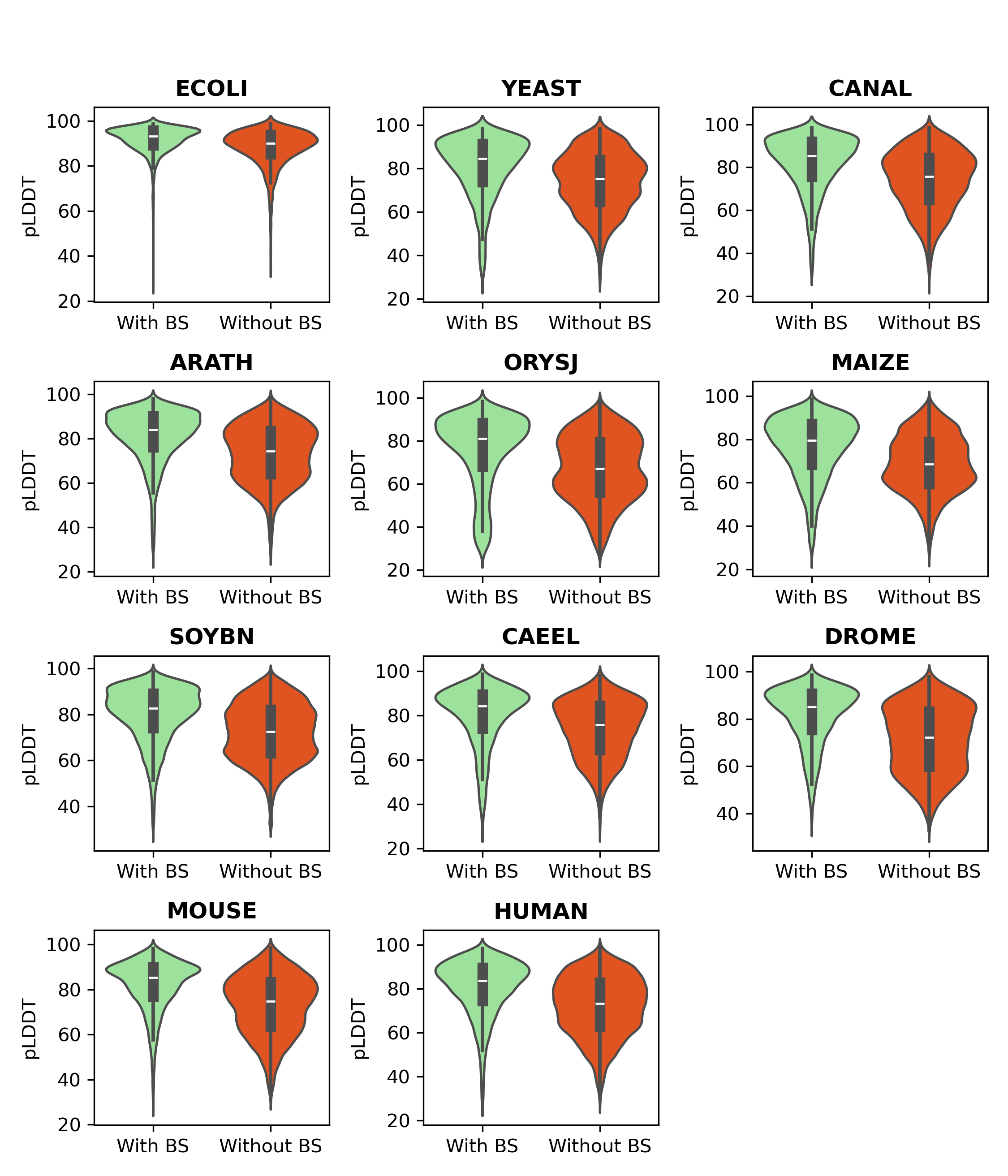

### S2Fig.tiff

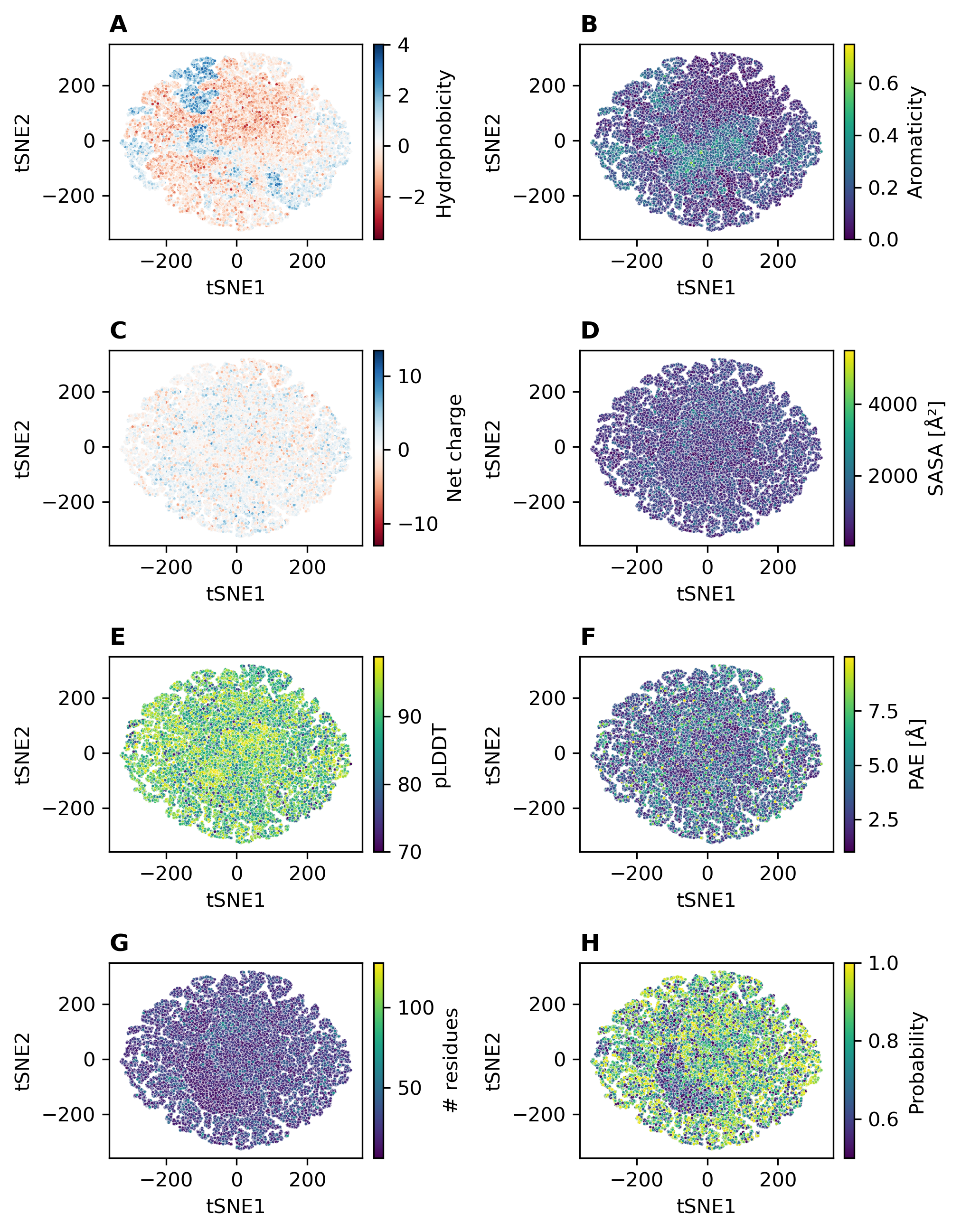

### S3Fig.tiff

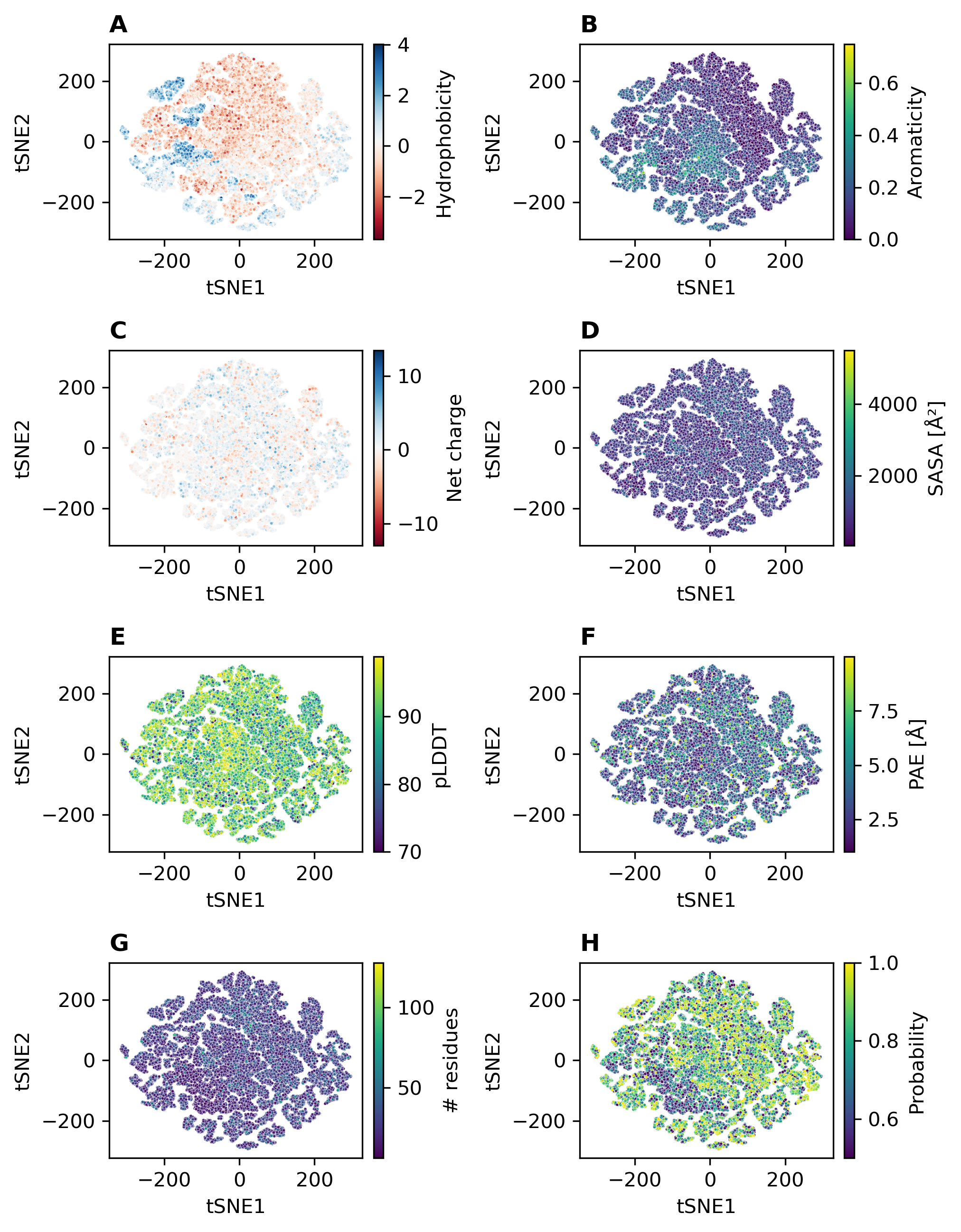

### S4Fig.tiff

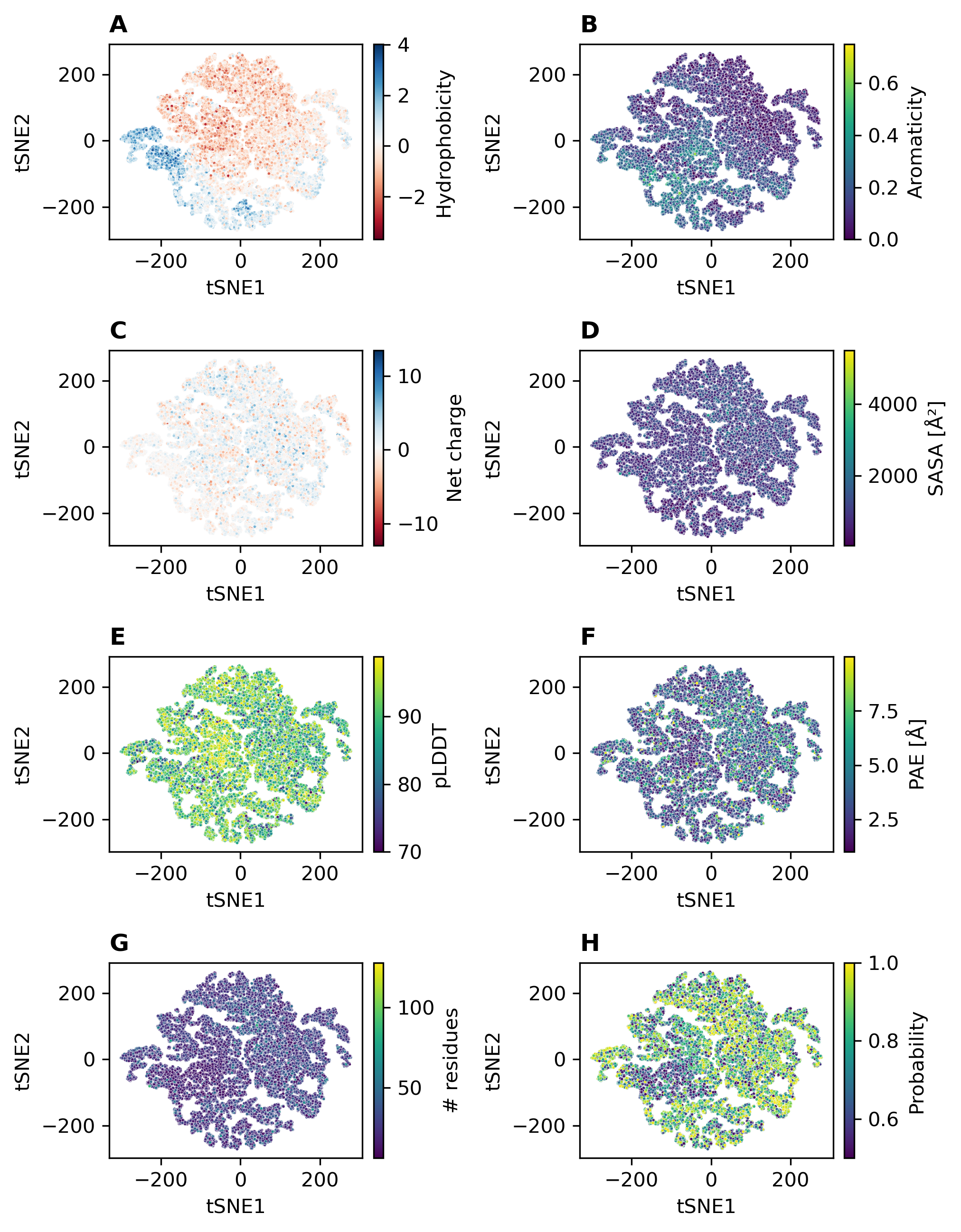

### S5Fig.tiff

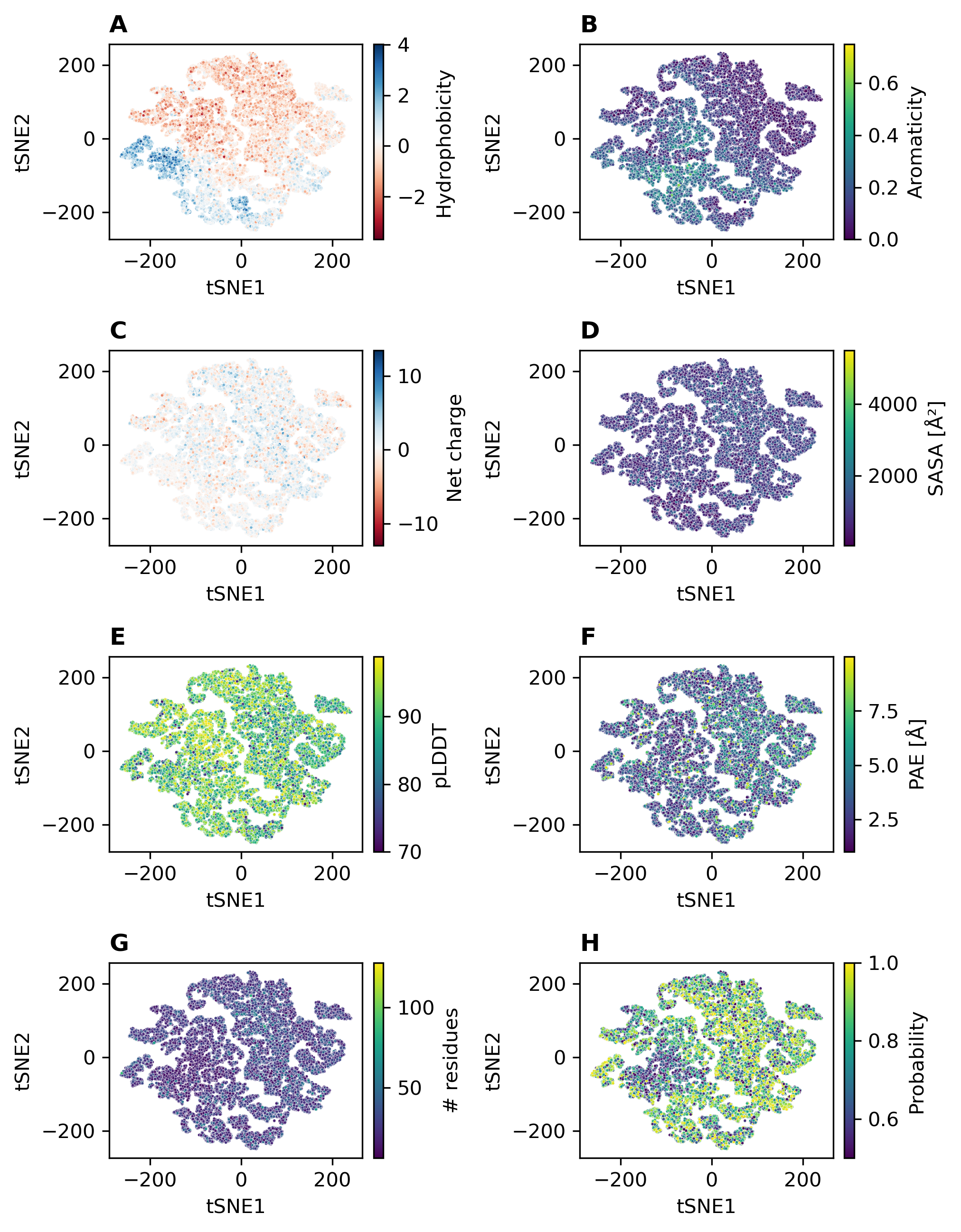

### S6Fig.tiff

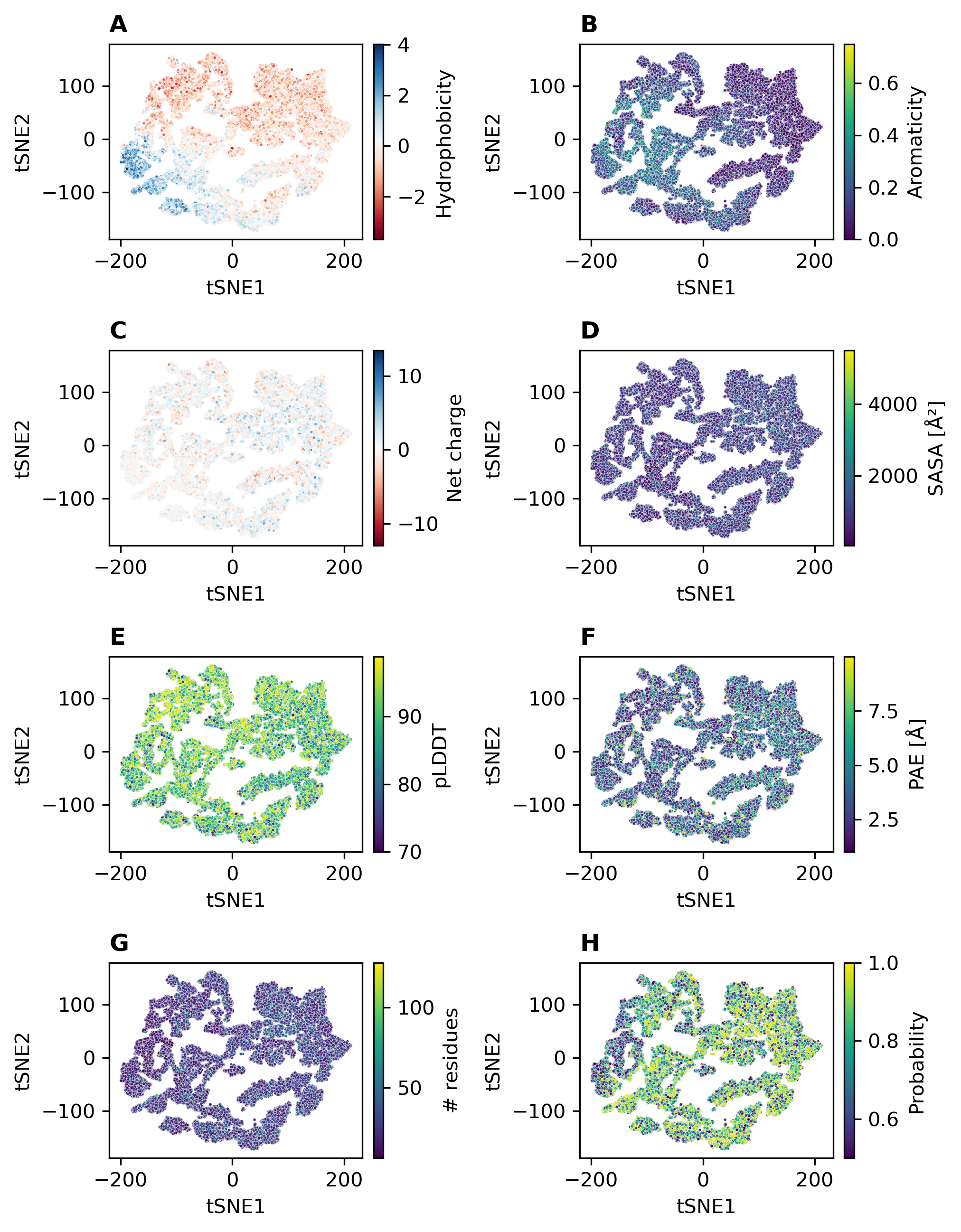

### S7Fig.tiff

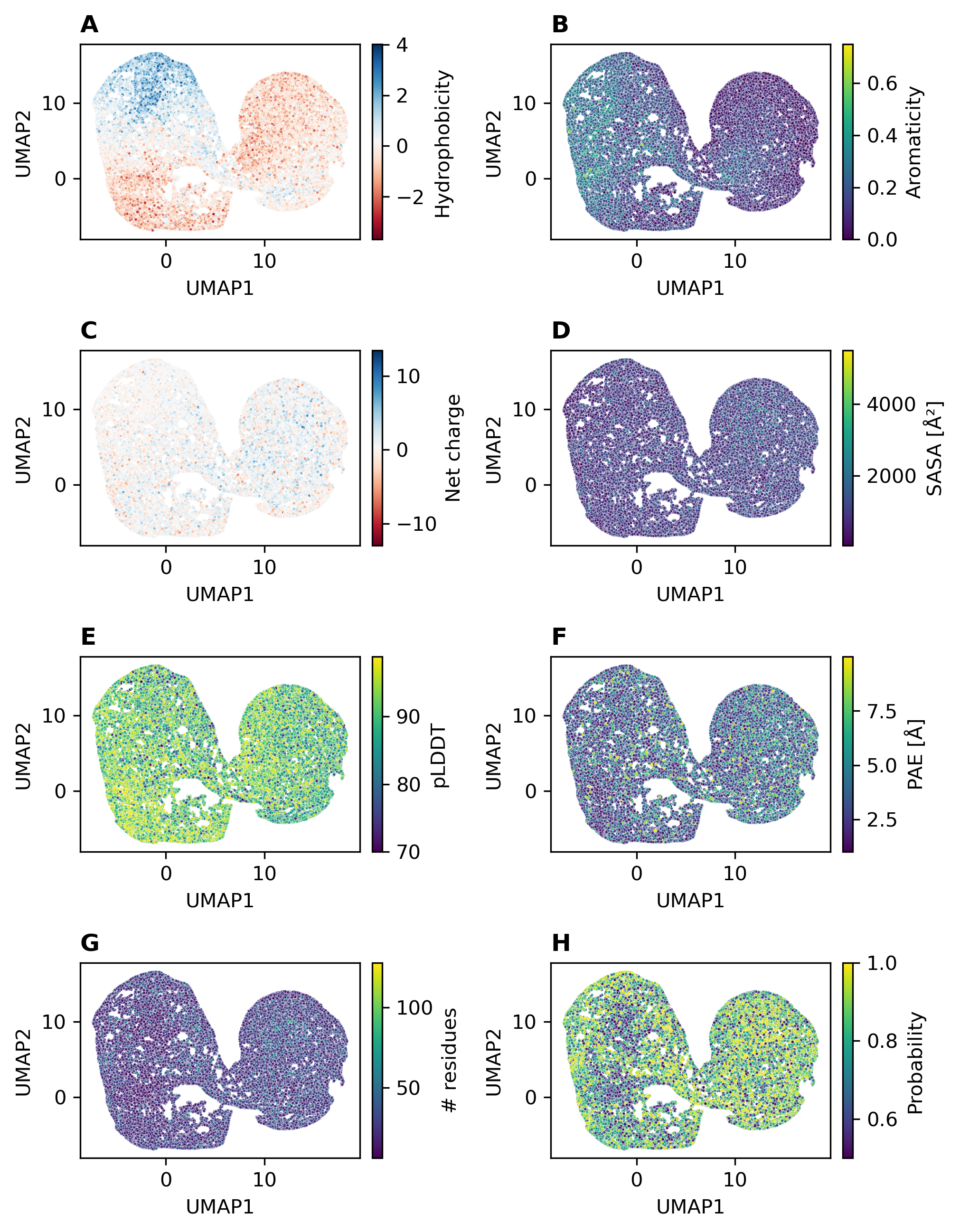

### S8Fig.tiff

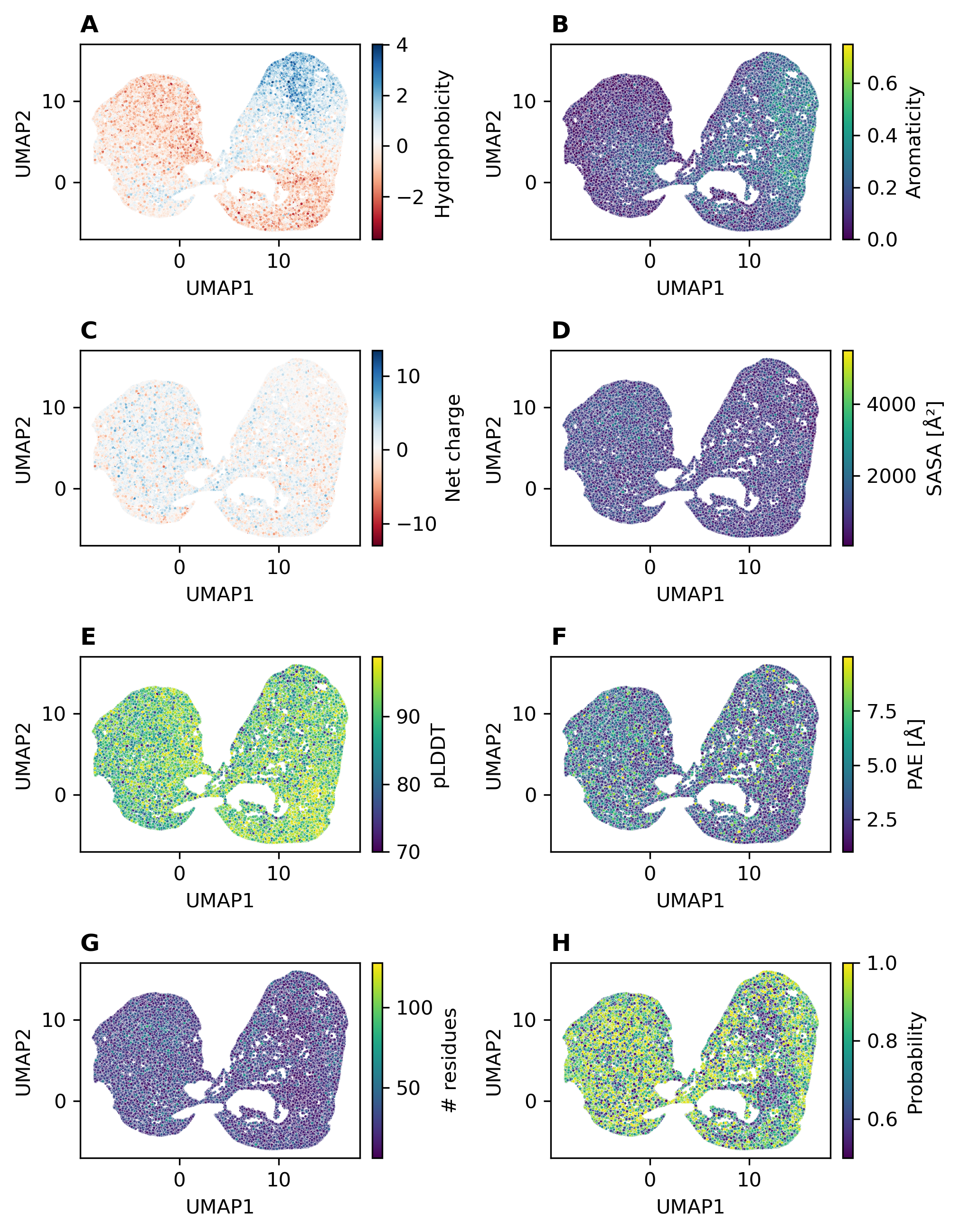

### S9Fig.tiff

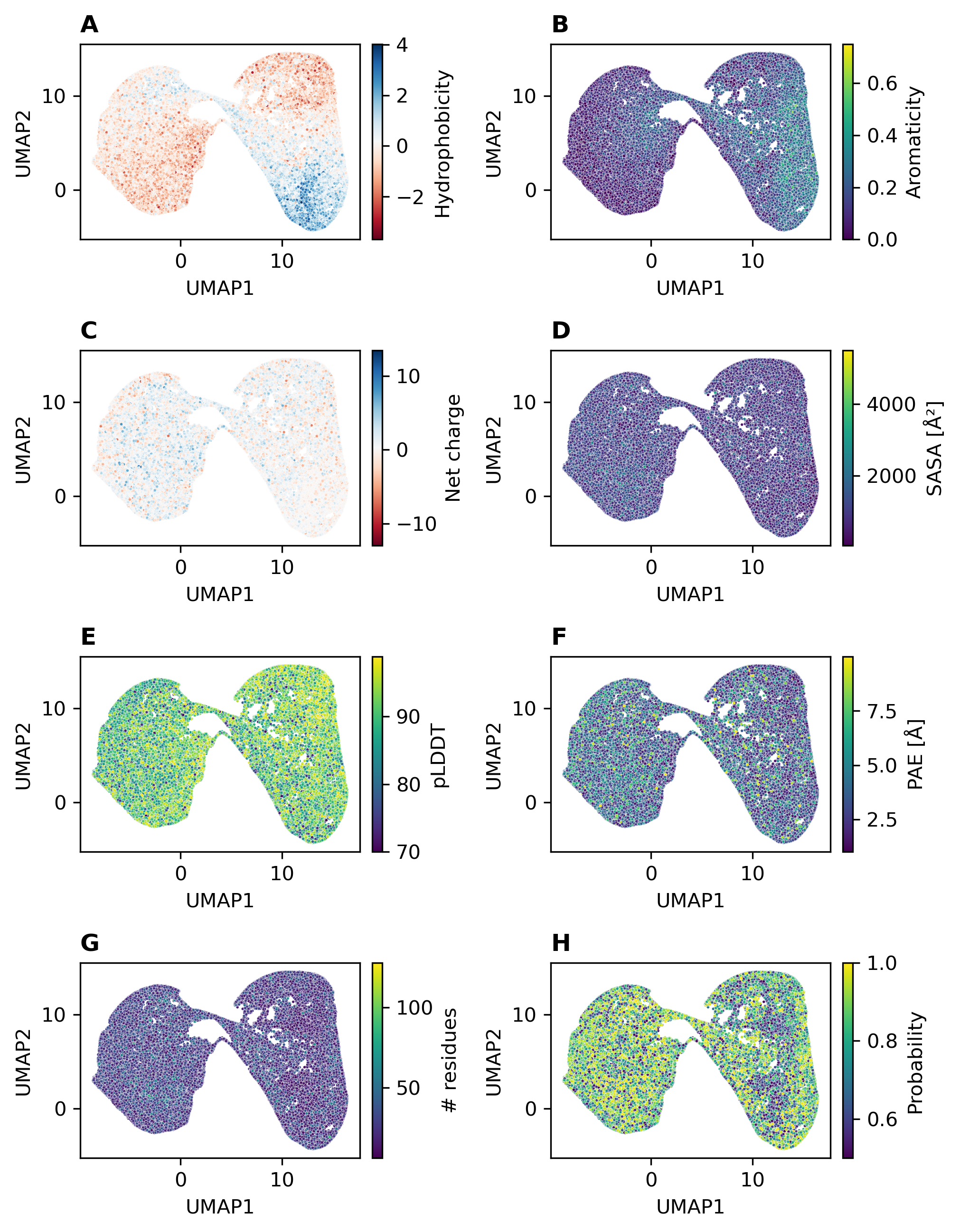

### S10Fig.tiff

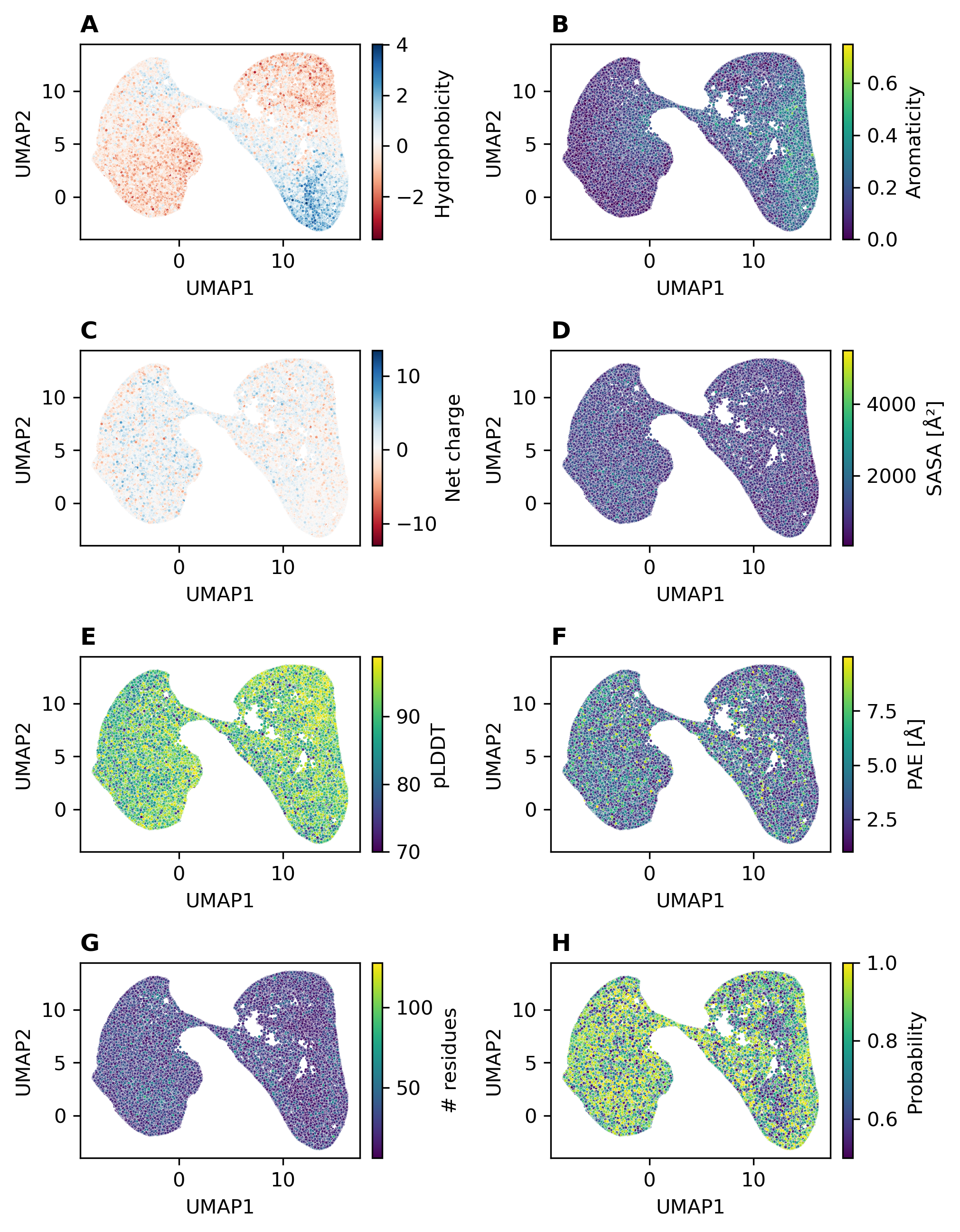

### S11Fig.tiff

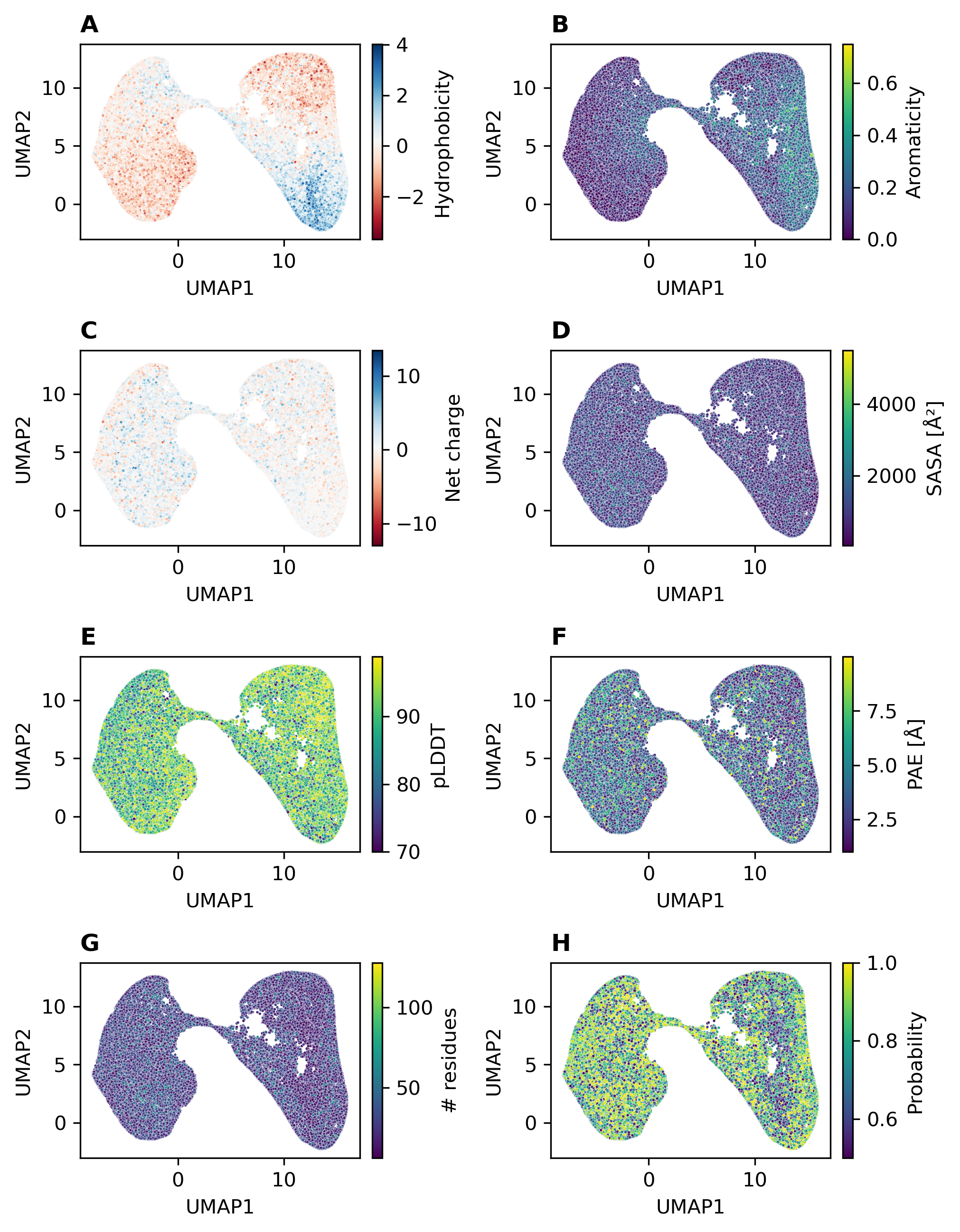

### S12Fig.tiff

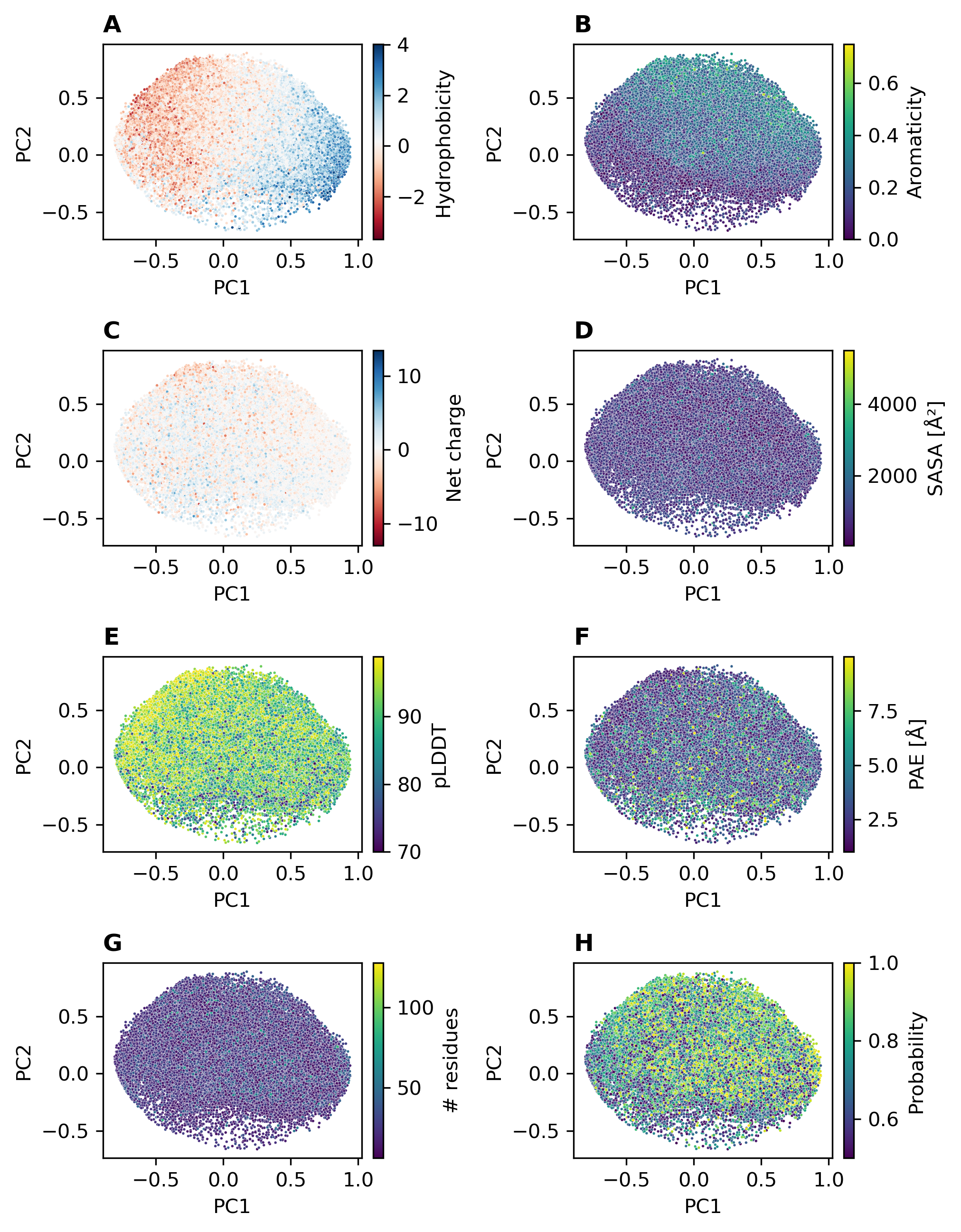

### S13Fig.tiff

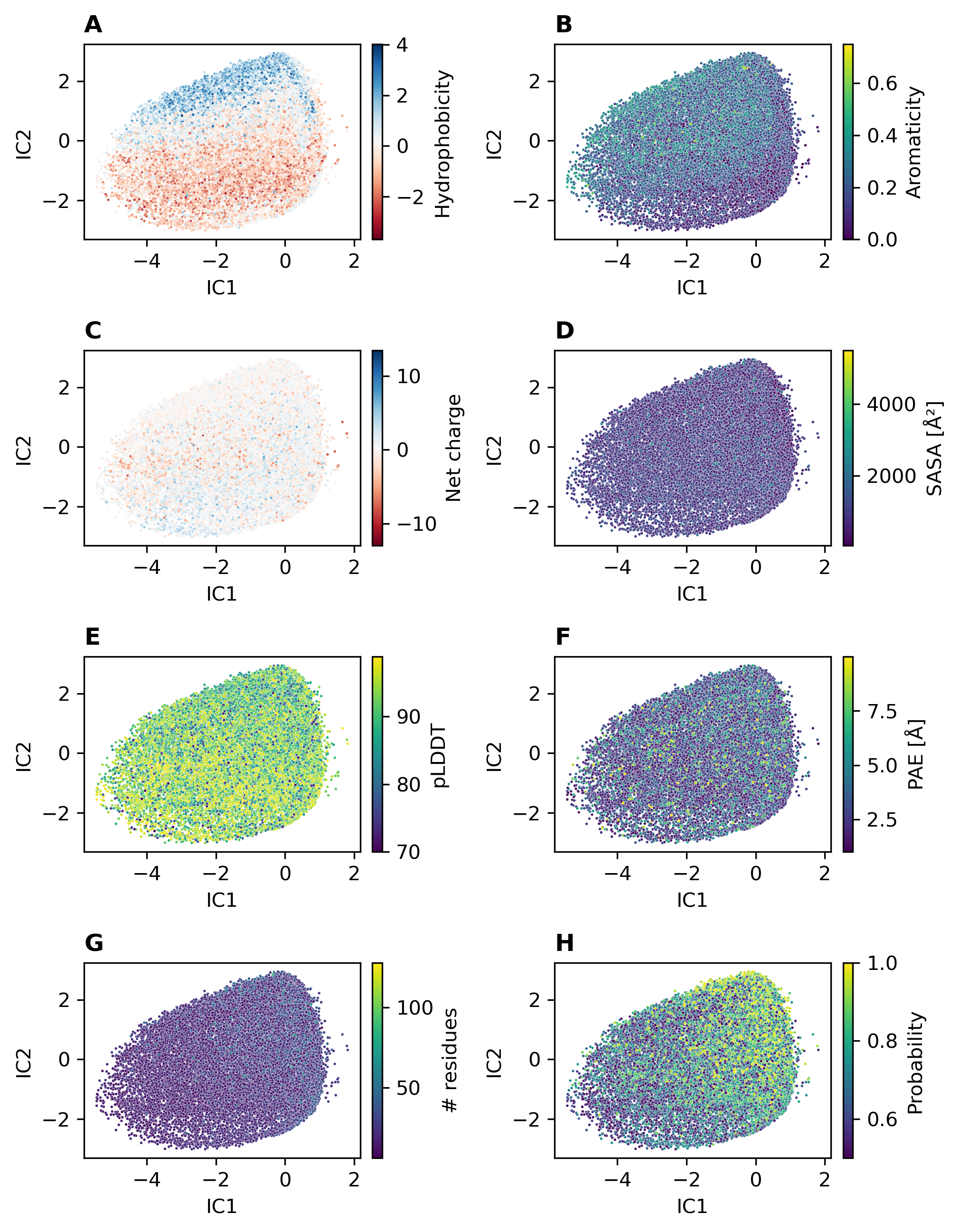

### S14Fig.tiff

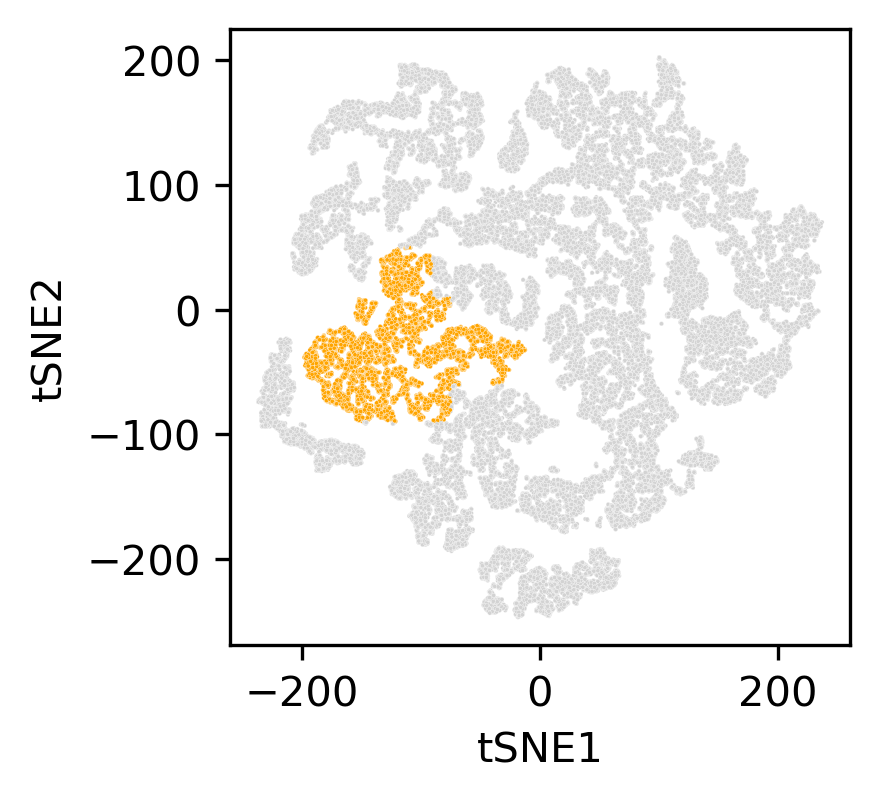

### S15Fig.tiff

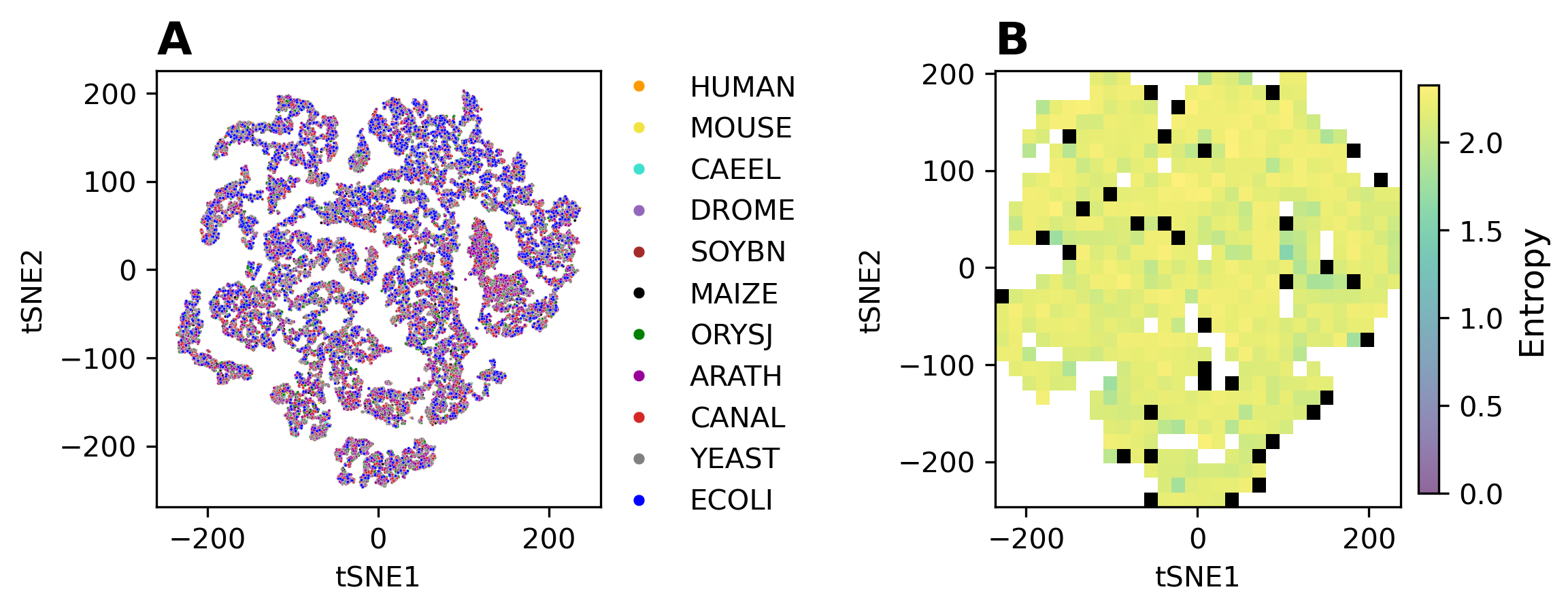
